# A small molecule probe elucidates the role of mitochondrial translocase TIMM44 in PINK1/Parkin regulated mitophagy

**DOI:** 10.1101/2022.02.16.480676

**Authors:** Michael A. Conti, Eric Torres, Alexander M. van der Bliek, Carla M. Koehler

**Affiliations:** Department of Chemistry and Biochemistry, University of California, Los Angeles, CA 90095, USA; Biological Chemistry, University of California, Los Angeles, CA 90095, USA; Molecular Biology Institute, University of California, Los Angeles, CA 90095, USA; Jonsson Comprehensive Cancer Center, University of California, Los Angeles, CA 90095, USA

**Keywords:** Mitochondria, small molecule probe, protein translocation, chemical biology, mitophagy, translocase of mitochondrial inner membrane, mitochondrial protein import

## Abstract

Neurodegenerative diseases have been linked to a dysfunctional mitochondrial quality control system that is partially maintained by proteins PINK1 and Parkin. Whereas mitophagy pathways are becoming well-characterized, less is known about the molecular mechanisms of PINK1 trafficking in mitochondria. Accordingly, we have used a small molecule probe (MitoBloCK-10/MB-10) that modulates the activity of TIMM44, an essential component of the protein associated motor (PAM) complex for the mitochondrial inner membrane (TIM23) translocase, to characterize PINK1 import. MB-10 did not inhibit the import or degradation of PINK1 in energized mitochondria. However, when mitophagy was induced by the addition of an uncoupler or respiratory inhibitor, MB-10 treatment altered PINK1 trafficking by inhibiting association with the TOM complex and impairing Parkin recruitment and subsequent mitophagy. MB-10 analogs that did not inhibit TIMM44 activity failed to impair mitophagy, thereby assigning specificity to MB-10. Because PINK1 undergoes lateral release from the TIM23 translocon to interact with inner membrane (IM) modulators, our studies support that TIMM44 may be a key regulator and that the PAM complex has a central role in regulating PINK1-dependent mitophagy. Our studies also provide a probe for dissecting PINK1/Parkin events for mitochondria as well as studying PINK1-dependent mitophagy in cell and animal models.

## Introduction

Deregulation of mitophagy, the process of clearing dysfunctional mitochondria, has been implicated in various human diseases, including neurodegeneration [1]. Certain mutations in PTEN-induced putative kinase (PINK1), a mitochondrial serine kinase, and Parkin, a cytosolic E3-ubiquitin ligase, lead to early onset familial Parkinson’s disease (PD) [2]. These proteins initiate the selective turnover of damaged mitochondria [3], which can be induced by the loss of the mitochondrial membrane potential [4] or accumulation of misfolded proteins in the mitochondrial matrix [5]. PINK1 and Parkin coordinate the ubiquitination of outer membrane (OM) proteins for proteasomal degradation [6,7] and the recruitment of autophagic machinery for eventual mitophagy [8–10].

General events of this pathway are well established. In healthy mitochondria, PINK1 is constitutively targeted to the mitochondria by a short N-terminal mitochondrial targeting sequence (MTS) of 34 amino acids [11,12], and is partially imported through the Translocase of the Outer Membrane (TOM) and the Translocase of the Inner Membrane (TIM23) complexes. The matrix processing peptidase (MPP) removes the PINK1 MTS [13] and PARL cleaves PINK1 between Ala103 and Phe104 in the IM [14]. This processed form of PINK1 undergoes proteasomal degradation, leading to rapid turnover of the PINK1 pool in healthy mitochondria [15]. Upon mitochondrial insult caused by treatment with uncouplers, the MTS of PINK1 is not cleaved, causing the accumulation of the precursor form at the OM with the kinase domain remaining in the cytosol [16]. PINK1 at the OM has the dual activity of phosphorylating Parkin at Ser65 and ubiquitin at Ser65 [17–19], both of which activate the ligase function of Parkin to maximize ubiquitination activity at the mitochondrion [20]. This pathway not only stimulates ubiquitin-proteasome degradation of select OM proteins, but also recruits and activates the autophagy machinery at mitochondria [10].

Conversely, the molecular mechanisms for PINK1 translocation into mitochondria are complex. It has been shown that TOMM40 is important for the import of precursor PINK1 and the association of PINK1 with the OM [16]. TOMM7, a homolog of the *S. cerevisiae* protein Tom7 that modulates TOM complex dynamics and functions in protein import [21], is necessary for PINK1 accumulation on the OM but not for the PARL cleavage of PINK1 [22]. Additionally, import studies have revealed that PINK1 contains a second targeting sequence, referred to as the outer mitochondrial membrane localization signal (OMS), between the canonical MTS (amino acids 1-34) and the transmembrane domain at amino acids 94-110 [12]. With energized mitochondria, the MTS reaches the matrix to initiate the PINK1 degradation pathway, and the OMS remains dormant. But upon depolarization, the MTS does not engage the IM and the OMS becomes functional to associate with the TOM complex and activate the mitophagy pathway [12]. A recent study by Youle and colleagues added another layer of regulation [23]. PINK1 is released laterally to the IM when TIMM23 was reduced by RNA interference or TOM7 was deleted. PINK1 was subsequently cleaved by the IM protease OMA1 and then degraded by the proteasome. Thus, events at both the OM and IM seem to regulate PINK1 import dynamics.

Characterization of the PINK1 translocation pathway into healthy mitochondria and its retention on the OM in dysfunctional mitochondria may facilitate the development of therapeutic strategies [24]. Although approaches such as RNAi to study the PINK1 import pathway have been used and genetic screens have identified regulators of Parkin [22,25], there remains limited techniques to dissect molecular events of PINK1 at the mitochondria or to recapitulate the pathway in a specific, controllable manner. Currently, investigators commonly utilize the protonophore carbonyl cyanide m-chlorophenyl hydrazone (CCCP) or the expression of misfolded proteins targeted to the matrix to activate PINK1/Parkin-mediated mitophagy [5,26], but more precise methods are needed.

Here we report the characterization of a small molecule probe in PINK1/Parkin dynamics. This probe was identified in a chemical screen for mitochondrial protein import inhibitors in yeast. A Tim44 modulator, MitoBloCK-10 (MB-10) [27], had no effect on PINK1 import into healthy mitochondria, but inhibited the accumulation of PINK1 on the mitochondrial OM in the presence of CCCP or respiratory inhibitors, reducing the progression of mitophagy. These molecular studies further elucidate the mechanism of PINK1 import, specifically the role of TIMM44, and should provide a valuable tool for studies in model systems.

## Results

### MB-10 targets TIMM44 and inhibits PINK1 accumulation on the OM but not import

Because MB-10 (Figure S1A) binds to yeast Tim44, we first confirmed that human and yeast homologs were conserved. Sequence analysis shows that four of the seven Tim44 residues critical for MB-10 binding are identical in TIMM44, with a fifth residue sharing similarity (Figure S1B). Prior studies have also shown that the binding region of MB-10 shares structural similarities in yeast and human [Josyula, 2006 #3466]. Furthermore, TIMM44 was localized to the matrix and is loosely integrated in the IM, most likely in a protein-rich environment, similar to yeast Tim44 (Figure S1C,D). As in yeast mitochondria, MB-10 only inhibited the import of matrix-targeted precursors and not OM (MFN-1) or intermembrane space (IMS; CHCHD2) precursors (Figure S2A-F). Based on these results and a previous study [27], we confirm that MB-10 inhibits TIMM44 activity, likely by holding the TIM23 precursor tightly in the translocon.

PINK1 is targeted to the mitochondrion via its MTS and uses the TOM and TIM23 translocons for import, but arrests during translocation with the cytosolic presentation of its kinase domain at the OM. Import studies with radiolabeled Pink1 followed by trypsin sensitivity confirmed this localization (Figure S1E) [16]. At increased trypsin concentration, full-length PINK1 (designated FL-PINK1) was degraded, whereas control Su9-DHFR in the mitochondrial matrix was protected.

We investigated the role of the PAM complex in PINK1 import and stability by inhibiting TIMM44 activity with MB-10 treatment (Figure 1A). When HeLa cells (lacking Parkin expression) were treated with 10 μM CCCP to uncouple mitochondria, PINK1 (designated full-length PINK1, FL-PINK1) accumulated on the OM. In mitochondria pretreated with increasing concentrations of MB-10 (up to 40 μM) followed by CCCP addition, PINK1 accumulation on the mitochondrial OM was significantly reduced (Figure 1A,B). In contrast, treatment with MB-10 alone (in the presence of vehicle control DMSO) did not alter the accumulation of PINK1 on mitochondria (Figure 1A,B). As controls, proteasome inhibitor MG132 caused the accumulation of PARL-cleaved PINK1 (PINK1*) and valinomycin, a potassium ionophore, resulted in acute accumulation of full length PINK1. The OM protein MFN-1 is a typical substrate of Parkin that is ubiquitinated. However, MFN-1 was not ubiquitinated because the HeLa cells do not express Parkin.

**Figure 1.**
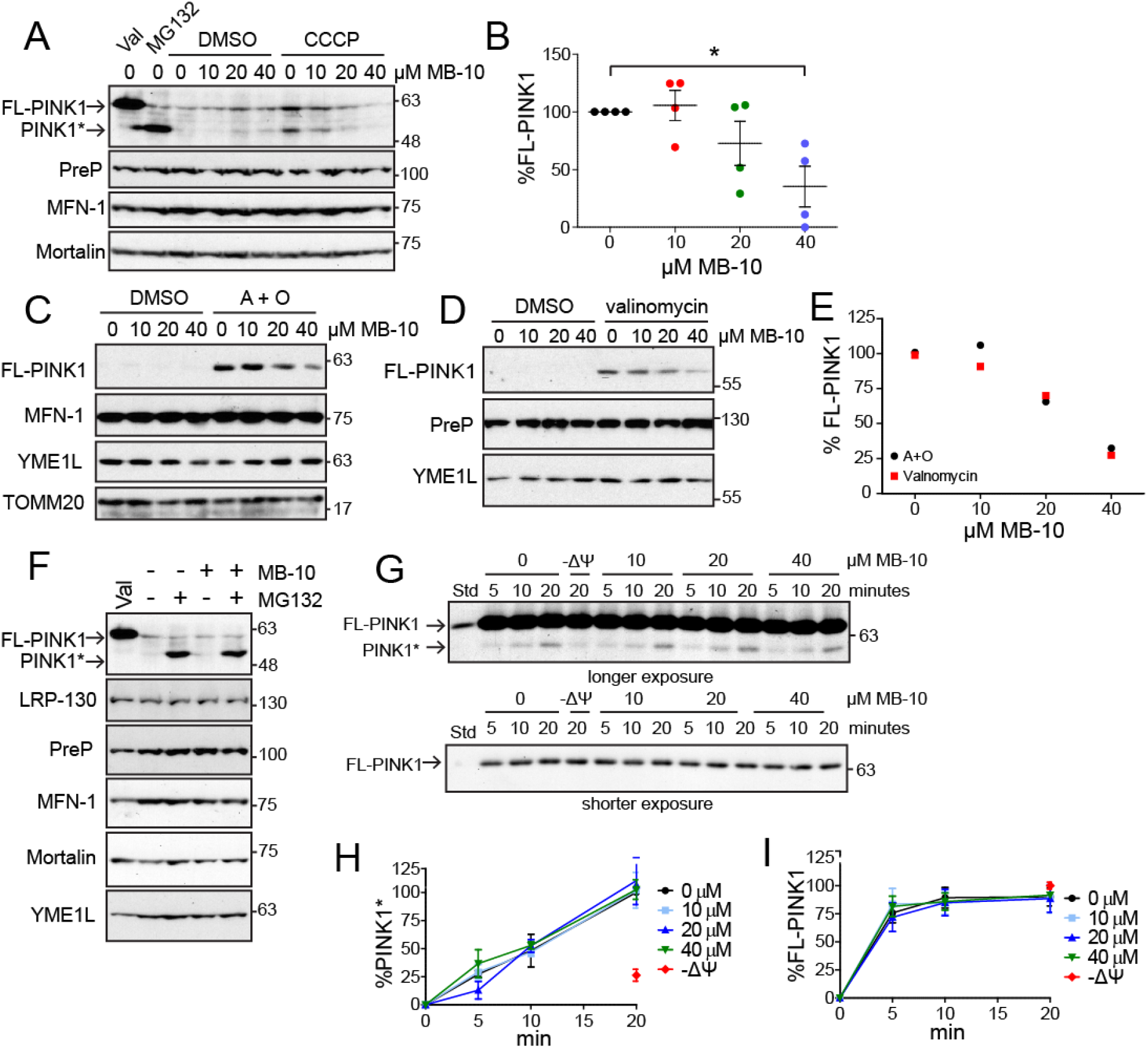
MB-10 inhibits the accumulation of PINK1 on compromised mitochondria, but does not affect PINK1 import into healthy mitochondria. **A.** HeLa cells were treated with the indicated concentration of MB-10 for 2 hrs, followed by the addition of 1 μg/mL valinomycin, 1% DMSO, 10 μM MG132, or 10 μM CCCP for an additional 3 hrs as indicated. The mitochondrial fraction was separated by SDS-PAGE, and the indicated proteins were detected by immunoblot. FL-PINK1, full-length PINK1; PINK1*, PARL-cleaved PINK1. **B.** Quantification of the FL-PINK1 band from ‘A’ under conditions in which the cells were treated with MB-10 and CCCP. The band at 0 μM MB-10 with CCCP represents 100% for each trial. The data represent the average ± SD of *n* = 4 trials. \**P* < 0.05 marks the difference between 0 and 40 μM MB-10. **C.** HeLa cells were treated as in ‘A’ with 10 μM antimycin A and 1 μM oligomycin (A + O) instead of CCCP. **D.** HeLa cells were treated as in ‘A’ with 1 μg/mL valinomycin for 1 hr instead of CCCP. **E. Quantification of the PINK1 band from ‘C’ and ‘D’** under conditions in which the cells were treated with MB-10 and CCCP. The band at 0 μM MB-10 with CCCP represents 100%. **F.** HeLa cells were treated with DMSO or 40 μM MB-10 for 2 hrs, followed by the addition of DMSO, 1 μg/mL valinomycin, or 10 μM MG132 for an additional 3 hrs. Samples were processed as in ‘A’. **G.** Upper—a representative image of an import assay with [^35^S]-PINK1 into isolated HeLa mitochondria that were pretreated with MB-10 or DMSO for 15 min. As an import control, mitochondria were uncoupled (-Δψ) with 50 μM CCCP. The longer exposure shows the PARL-cleaved form of PINK1, designated PINK*. Lower—a shorter exposure showing full-length PINK1, designated FL-PINK1. **H.** Quantification of the PARL-cleaved band PINK1* from ‘F’. The band at 0 μM MB-10 at 20 min represents 100% for each trial. The data represent the average ± SD of *n* = 3 trials. **I**. As in ‘G’ for FL-PINK1 from the shorter exposure; the −Δψ band represents 100% for each trial.

Loading controls (Mortalin and PreP) indicated that mitochondrial proteins were stable. We also tested the combination of antimycin A (complex III inhibitor) and oligomycin (ATP synthase inhibitor) or valinomycin as in Figure 1A. Similarly, PINK1 accumulation on the OM was decreased by MB-10 treatment with these additional uncouplers (Figure 1C-E). Thus, MB-10 treatment blocked PINK1 accumulation on the OM when the mitochondrial membrane potential was compromised with various inhibitors.

To determine if the reduction in PINK1 on mitochondria was caused by inhibiting the initial import of PINK1 into mitochondria, cells were treated with both MB-10 and MG132 followed by analysis of PINK1 localization on mitochondria (Figure 1F). In the presence of MG132, the accumulation of PARL-cleaved PINK1 (PINK1*) was identical in the presence of DMSO or MB-10. Import studies with radiolabeled PINK1 in the presence of MB-10 were similar to those with cells in Figure 1F (Figure 1G-I). Specifically, MB-10 treatment did not alter PARL cleavage of imported PINK1 (PINK1*) into isolated HeLa mitochondria, as shown in the autoradiograph with a longer exposure (Figure 1G). In contrast, uncoupler addition resulted in loss of the PARL-cleaved PINK1 intermediate (Figure 1G,H), providing additional evidence that TIMM44 does not play a direct role in PINK1 import in coupled mitochondria. In addition, the association of full-length PINK1 with mitochondria did not change in the presence of MB-10, as shown in the autoradiograph with a shorter exposure (Figure 1G,I†). Thus, MB-10 treatment did not alter normal PINK1 import in energized mitochondria.

### MB-10 treatment inhibits Parkin recruitment and mitophagy in uncoupled mitochondria

Blue-native gel analysis has been used to show that PINK1 associates with the TOM complex in uncoupled mitochondria [16]. We adapted this approach and confirmed that radiolabeled PINK1 imported into isolated HeLa mitochondria associated with the TOM complex in the presence of uncoupler, CCCP (Figure 2A). However, pretreatment with MB-10 followed by import in the presence of uncoupler resulted in a decreased amount of PINK1 at the TOM complex. In coupled mitochondria, the association of PINK1 with the TOM complex in the presence of DMSO or MB-10 alone was similar, confirming that MB-10 treatment did not alter PINK1 trafficking in coupled mitochondria.

**Figure 2.**
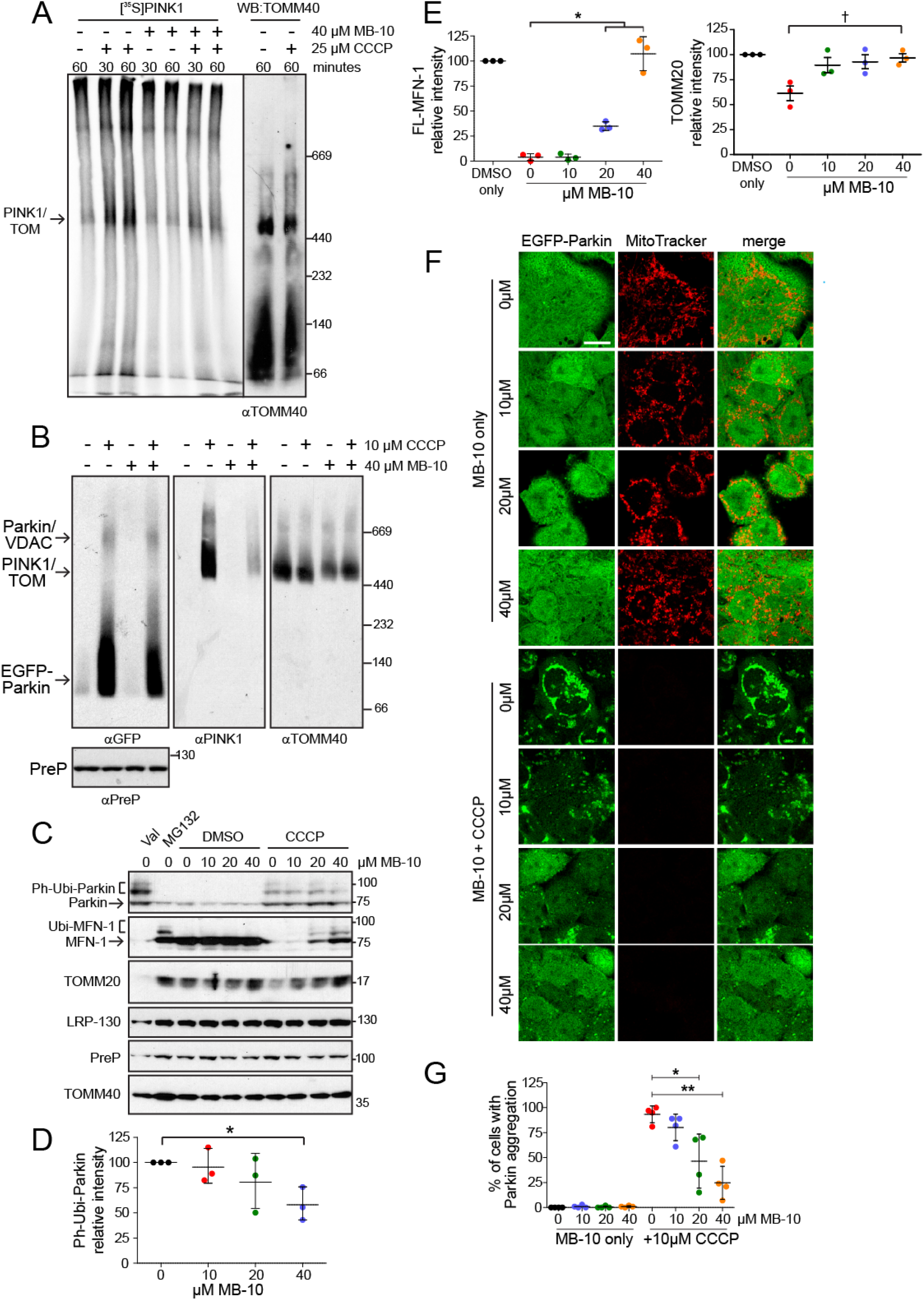
MB-10 disrupts association of PINK1 with the TOM complex and inhibits CCCP-induced aggreagation of Parkin. **A.** Radiolabeled PINK1 was imported into isolated HeLa cell mitochondria in the presence of MB-10 and/or CCCP as indicated. Mitochondria were lysed with 1% digitonin and lysates were separated on BN-PAGE. (left panel, [^35^S]PINK1; right panel, TOMM40 immunoblot from the same gel). **B.** HeLa cells overexpressing integrated EGFP-Parkin were treated with MB-10 for 2 hrs followed by the addition of CCCP for another 3 hrs. Samples were analyzed by BN-PAGE as in ‘A’ followed by immunoblot. PreP is included as a loading control from the same samples run on SDS-PAGE. **C.** HeLa cells with EGFP-Parkin were treated as in Figure 1A. Ph-Ubi-Parkin marks the phospho-ubiquitinated form and Ubi-MFN-1 marks the ubiquitinated form. LRP-130, PreP, and TOMM40 were included as loading controls. **D.** Quantification of the Ph-Ubi-Parkin band. The band at 0 μM MB-10 with CCCP represents 100% for each trial. The data represent the average ± SD of *n* = 3 trials (\**P* < 0.05). **E.** Quantification of the FL-MFN-1 band and the TOMM20 band. For each, the band at DMSO only represents 100% for each trial. The data represent the average ± SD of *n* = 3 trials (\**P* < 0.05; †*P*<0.1). **F.** Representative fluorescent images after treatment with indicated concentrations of MB-10 for 2 hrs, followed by the addition of either DMSO of 10 μM CCCP for an additional 3 hrs. Scale bar = 15 μm. **G.** Quantification of Parkin aggregation in ‘F’ by cell counting. 100 cells were counted for each condition; data represent the average ± SD of *n* = 4 trials (\**P* < 0.05).

We also used HeLa cells overexpressing EGFP-Parkin to determine if subsequent steps in mitophagy were impaired [8]. HeLa cells expressing EGFP-Parkin were pretreated with MB-10 or DMSO followed by CCCP addition. As in Figure 2A, PINK1 association with the TOM complex was increased in cells treated with CCCP (Figure 2B). Parkin recruitment to mitochondria, including the Parkin-VDAC complex [28], was also detected. However, pretreatment with MB-10 followed by uncoupler addition resulted in decreased PINK1 association with the TOM complex in as well as a slight reduction in the association of Parkin with mitochondria (Figure 2B). To confirm Parkin association with the mitochondria, the mitochondrial fraction was tested by SDS-PAGE (Figure 2C). Because Parkin expression in these cells is high, a low level of Parkin was associated with mitochondria even in untreated cells (Figure 2C). However, an increase in phospho-ubiquitinated Parkin (designated Ph-Ubi-Parkin) on mitochondria was detected following CCCP treatment in the absence of MB-10 (Figure 2C,D). Pretreatment with an increasing concentration of MB-10 significantly decreased association of phospho-ubiquitinated Parkin in the presence of CCCP (Figure 2C,D). OM proteins are ubiquitinated by Parkin and then degraded by the proteasome during mitophagy, and MFN-1 is a useful OM marker of this processbecause it is ubiquitinated and degraded [7]. In correlation with ubiquitinated Parkin, MFN-1 was degraded and the abundance of TOMM20 was also decreased (Figure 2C-E). However, MB-10 treatment significantly reduced the amount of phospho-ubiquitinated Parkin (Figure 2C,D) and the subsequent degradation of MFN1 and TOMM20 (Figure 2C,E) in cells treated with uncoupler.

We also tracked the distribution of EGFP-Parkin within cells treated with MB-10 by fluorescent microscopy (Figure 2F,G). Under normal conditions (DMSO), Parkin localization was diffuse throughout the cytosol; and the mitochondrial network, as marked by robust MitoTracker staining, was distributed throughout the cell. With an increasing MB-10 concentration, Parkin localization remained diffuse and the membrane potential was maintained. However, the mitochondrial network fragmented with 40 μM MB-10, perhaps due to an alteration of dynamics proteins associated with the decrease in functional TIMM44 [29]. Upon addition of CCCP, the mitochondrial membrane potential was dissipated, causing strong Parkin aggregation. The addition of 20 and 40 μM MB-10 inhibited this Parkin aggregation even though membrane depolarization was induced by CCCP addition (Figure 2 F,G).

To study the kinetics of mitophagy in the presence of MB-10, we used the tandem sensor RFP-GFP-LC3. LC3 causes the sensor to localize to autophagic structures and the RFP and GFP fluorescence monitor autophagy progression [30], giving us a good proxy for mitophagy. In the acidic environment of the lysosome, the GFP is quenched, which results in an increase in red puncta from RFP and is indicative of mitophagy. In contrast, yellow/green puncta indicate that the LC3 has not entered autophagosomes. In MEF cells expressing Flag-Parkin [31] that were treated with CCCP alone, the number of autophagosomes increased in a 3-hour time course (Figure 3A,B). In addition, the amount of red puncta was greater than yellow/green puncta, indicating mitochondria were entering autophagosomes. Treatment with MB-10 significantly impeded mitophagy progression with a small increase in the number of autophagosomes after 3 hours (Figure 3A,B). Because PINK1/Parkin-mediated mitophagy can clear the entire population of mitochondria in 24 hrs [8,10], we tested if MB-10 treatment blocked mitophagy over longer periods by assessing the loss of mtDNA (Figure S3). CCCP treatment of the EGFP-Parkin overexpression cell line resulted in nearly complete removal of the mitochondria after 24 hrs as evidenced by a lack of mtDNA staining (in nucleoid puncta surrounding the cell nucleus) with an anti-DNA antibody and lack of mitochondria in MitoTracker treated cells. In contrast, MB-10 treatment inhibited the degradation of mtDNA and mitochondria in CCCP-treated cells, because mitochondrial DNA was detected and Parkin was localized at those puncta (Figure S3). Note that, at this much longer time point of 24 hrs, Parkin had been recruited to mitochondria in MB-10 treated cells, but mitochondria were still present. This data as well as results from Figure 2 showing that MB-10 inhibits Parkin recruitment at shorter time points indicate that MB-10 treatment slows clearance of mitochondria in cells in which mitophagy is induced.

**Figure 3.**
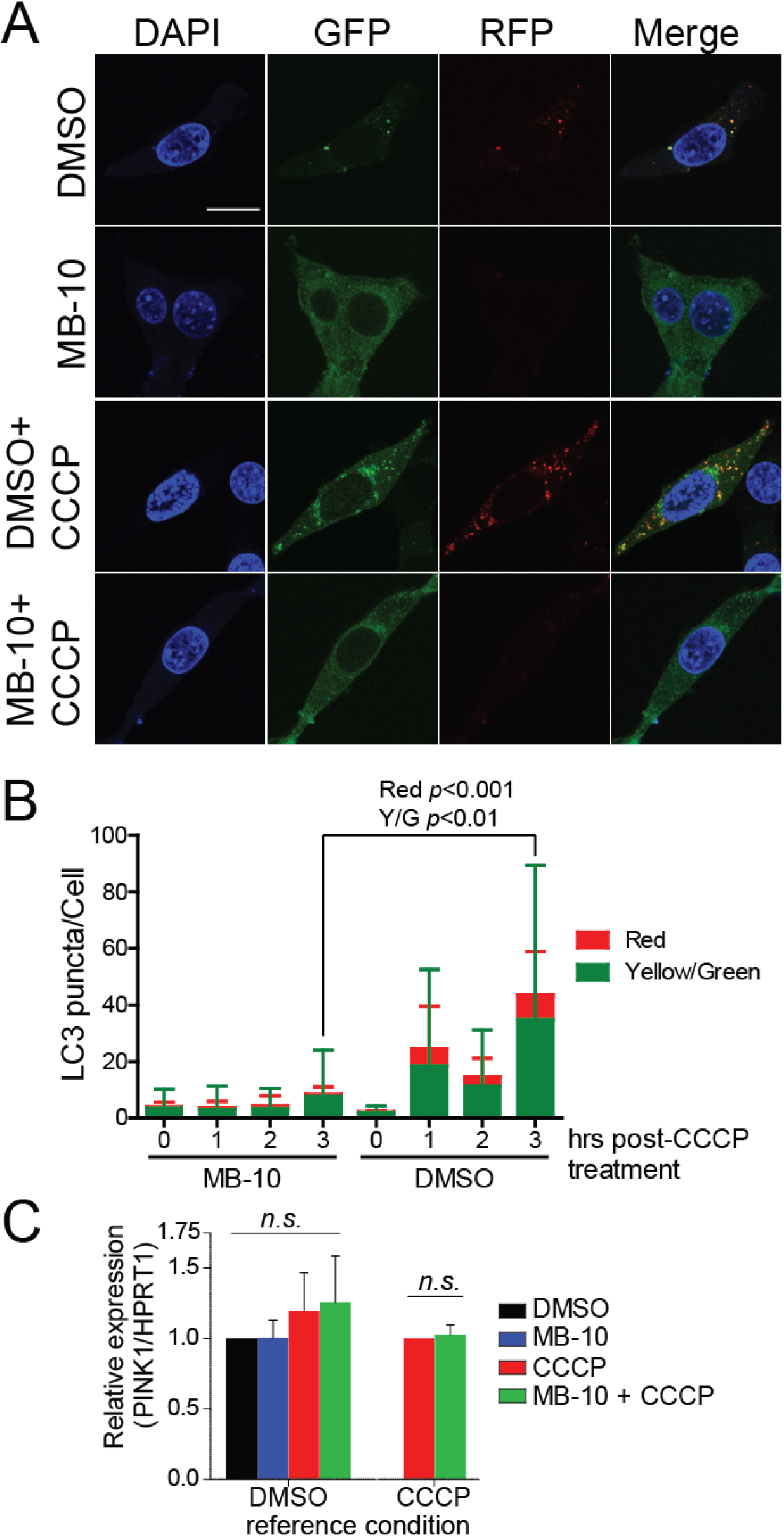
MB-10 treatment blocks autophagy, including mitochondrial, but does not affect PINK1 transcription. **A.** MEF cells stably expressing Flag-Parkin were transfected with LC3-EGFP-RFP. After 16 hrs, 40 μM MB-10 or 1% DMSO was added, followed by treatment with 10 μM CCCP. In a time course of 0 to 3 hours, cells were subsequently fixed in 4% paraformaldehyde and imaged at 63X. Representative images of the treated cells at 3 hrs are shown. Cells treated with MB-10 prior to CCCP treatment showed less autophagosome (green/yellow) and autolysosome (red) particles relative to cells treated with DMSO. Scale bar represents 15 μm. **B.** Quantification of 50 cells per conditions was performed using particle analysis of green/yellow and red particles. Data represent average ± SEM for 50 cells of *n* = 3 replicates. **C.** Analysis of *PINK1* expression by qPCR in HeLa cells treated with 10 μM CCCP and 40 μM MB-10 as indicated. Data represent average ± SEM of *n* = 3.

### Reduction in PINK1 accumulation by MB-10 is specifically to TIMM44 inhibition

Because MB-10 may have off-target effects, we addressed specificity using a variety of methods. Quantitative PCR (qPCR) analysis showed that *PINK1* expression in HeLa cells was not significantly different in the presence of MB-10 or CCCP (Figure 3C). Thus, the addition of the compounds is not inducing a nonspecific cellular stress that alters gene expression, which could reduce PINK1 protein levels.

We also used previously identified compounds from structure-activity relationship (SAR) studies, termed Analog-3 and −4, [27] that did not inhibit yeast Tim44. Analog-3 and −4 are structurally similar to MB-10 and are useful to address off-target effects of MB-10 (Figure 4A). These analogs did not inhibit the import of Su9-DHFR into isolated HeLa cell mitochondria, demonstrating that they do not inhibit TIMM44 function (Figure 4B,C). The compounds also did not disrupt the association of PINK1 with the TOM complex in the presence of CCCP (Figure 4D). Analysis of downstream events, including Parkin recruitment to mitochondria and ubiquitination and MFN-1 degradation, showed that these analogs do not block mitophagy (Figure 4E-H). Specifically, ubiquitinated Parkin and MFN-1 degradation were not decreased by treatment with Analog-3 and −4 (Figure 4E,F). With cells, aggregation of EFGP-Parkin was not inhibited in the presence of the Analog-3 and-4 when cells were treated with CCCP (Figure 4 G,H). Taken together, this data supports that the inhibition of PINK1 accumulation and block in mitophagy caused by MB-10 is not caused by off-target effects of the small molecule.

**Figure 4.**
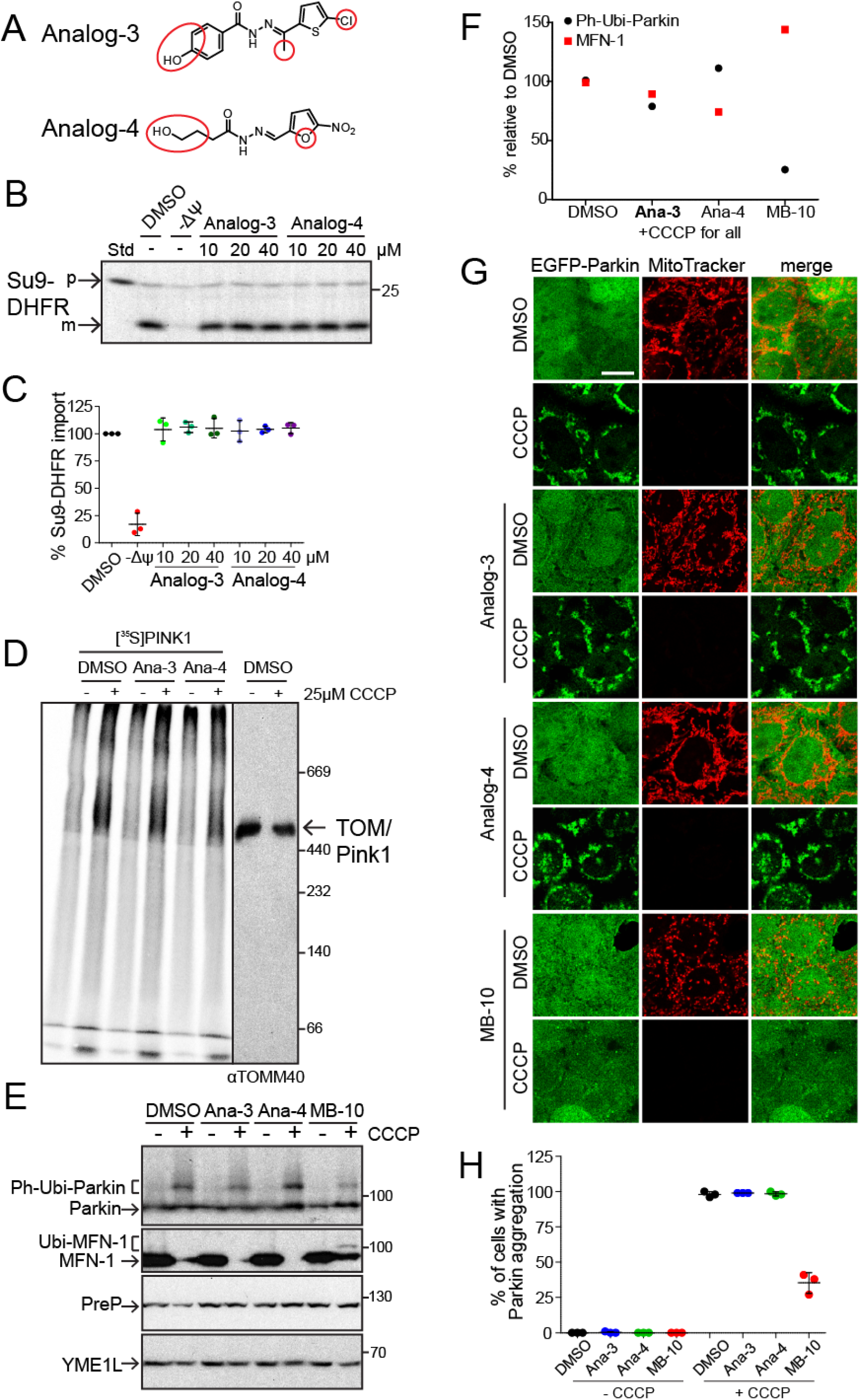
SAR compounds of MB-10 do not alter PINK1-dependent mitophagy. **A.** Two previously identified compounds (Analog-3 and −4) that have similar structures to MB-10 but do not inhibit TIMM44 activity were analyzed. Red circles indicate differences with the MB-10 structure. **B.** Representative image of an import with Su9-DHFR into isolated HeLa cell mitochondria that were pre-treated with Analog-3 or −4 for 15 min. p = precursor; m = mature form. **C.** Quantification of the mature band from ‘B’; the DMSO band represents 100%. Data represent average ± SD for *n* = 3 replicates. **D.** As in Figure 2A, Pink1 was imported into HeLa cell mitochondria for 60 min with Analog-3 and −4. **E.** As in Figure 2C, HeLa cells overexpressing EGFP-Parkin were treated with MB-10, Analog-3 or −4 followed by CCCP addition. Parkin and MFN-1 (and ubiquitinated forms) were detected. **F**. Quantification of Ph-Ubi-Parkin and MFN-1 from ‘E’ for the +CCCP condition for each molecule. Percentages are relative to the DMSO+CCCP condition. **G.** As in Figure 2F, representative fluorescent images of HeLa cells overexpressing EGFP-Parkin after treatment with MB-10, Analog-3, or −4 are shown. Scale bar = 15 μm. **H.** Quantification of Parkin aggregation in ‘F’ by cell counting. 100 cells were counted for each condition; data represent the average ± SD of *n* = 3 trials

In the absence of specific small molecule modulators, a typical approach is to use CRISPR or RNAi approaches to inactivate the protein of interest. However, a drawback to this approach is that RNAi takes several days and CRISPR may not be possible for proteins that are essential for viability. Because TIMM44 is essential, we used RNAi to knock-down the TIMM44 protein. The abundance of TIMM44 was decreased by approximately 95% with a marked impairment in the import of Su9-DHFR into isolated mitochondria (Figure 5A). However, the formation of the PARL-cleaved form of PINK1 in import and the association of PINK1 with the TOM complex were not decreased markedly as has been shown for MB-10 treatment (Figure 5B). Thus, the studies using RNAi in which TIMM44 is decreased failed to recapitulate those with MB-10. Using an RNAi or CRISPR approach to delete TIMM44 is not ideal because of the general impairment of mitochondrial function over a long time period and the potential adjustment of cells to reduced TIMM44 levels. This supports our approach with small molecules that rapidly impair TIMM44 function over a shortened period of 3-5 hours to study the impact on PINK1 trafficking and mitophagy in cell models.

**Figure 5.**
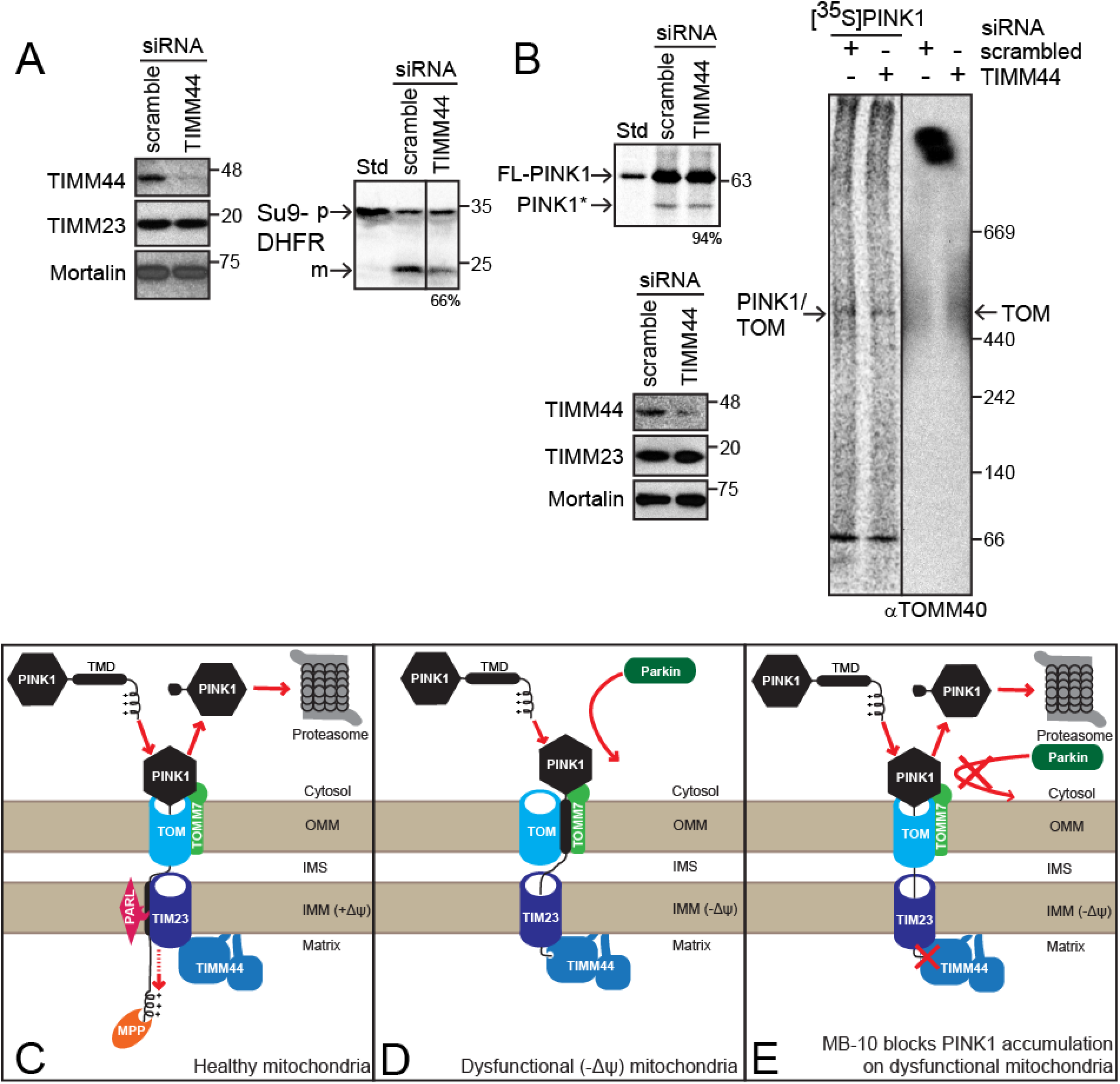
Pink1 trafficking is not impaired in cells with decreased TIMM44. **A.** HeLa cells were transfected with either scrambled or TIMM44 siRNA for 48 hours. TIMM44 knockdown was demonstrated by immunoblot analysis (left). Mitochondria were isolated followed by Su9-DHFR import; the percentage under the final panel represents the % import relative to scramble. **B.** HeLa cells were treated with siRNA as in ‘A’, followed by radiolabeled PINK1 import into the isolated mitochondria. Import reactions were analyzed on SDS-PAGE as in Figure 1F and BN-PAGE as in Fig 2A; for the SDS-PAGE import, the percentage under the final panel represents the % import relative to scramble. An immunoblot of the mitochondria is included to demonstrate TIMM44 knockdown. (C-E) Schematic depicting the activities of both TIMM44 and MB-10 **C.** PINK1 is partially imported into healthy mitochondria, cleaved by PARL, and degraded by the proteasome. TIMM44 plays no role. **D.** PINK1 accumulates on the OM on dysfunctional mitochondria (typically by uncoupler treatment) followed by Parkin recruitment and mitophagy. TIMM44 is a necessary component for PINK1 stability and therefore Parkin recruitment. **E.** Treatment with MB-10, a TIMM44 inhibitor, blocks the accumulation of PINK1 on damaged mitochondria in a form that precludes Parkin recruitment.

## Discussion

In this study, we have used a small molecule modulator of mitochondrial protein translocation to characterize PINK1/Parkin trafficking in mitochondria (Figure 5C-E). The data demonstrate that inhibiting TIMM44 with MB-10 treatment has no consequence on normal PINK1 import, cleavage, and ultimately degradation in healthy mitochondria (Figure 5C), which suggests that TIMM44 is not involved in normal PINK1 trafficking. Upon mitochondrial insult, inhibiting TIMM44 activity alters the trafficking of PINK1 at the mitochondrial OM and subsequently reduces Parkin recruitment (Figure 5D). This is unexpected because PINK1 is predicted to bypass the TIM23 translocon and TIMM44 in the absence of a membrane potential (Figure 5D). However, MB-10 studies support an important role for TIMM44 and the PAM motor for PINK1 presentation on the OM for subsequent Parkin recruitment. Previous studies showed that TOMM7 also lacked a direct role in PINK1 import, but was necessary for the association of PINK1 with the TOM complex [22], indicating that there may be coordination between the TOM and TIM23 complexes. Recent evidence also supports the notion that the TIM23 and PAM complex may not play a role in normal PINK1 import but are needed for PINK1 maintenance [23,32]. Because PINK1 contains two targeting sequences [12], it is possible that TIMM44 interacts exclusively with the second targeting sequence, which is critical for the interaction between PINK1 and the TOM complex.

### A role for TIMM44 and the PAM complex in PINK1 trafficking

A role for the PAM complex in PINK1 trafficking may seem unexpected, but recent studies support the importance of the TIM23 translocon, particularly the PAM complex. Interactions at the TIM23 translocon are critical for PINK1 trafficking. Under normal conditions, PINK1 may be cleaved by the IM protease PARL and then removed from the mitochondrion by retro-translocation and degradation by the proteasome [26,33]. This activity indicates that PINK1 must slide laterally out of the TIM23 translocon to reach PARL and that the PAM motor may be important for holding the N-terminus of PINK1 as it is transferred. A recent paper by Youle and colleagues [23] showed that PINK1 was imported into depolarized mitochondria of cells that lack TOMM7. Thus, a membrane potential was not essential for PINK1 to engage the TIM23 translocon. TIMM44 with its association in the membrane and the TIM23 channel may be positioned to tether PINK1 for lateral release from the translocon and subsequent cleavage by OMA1 and/or PARL under stress conditions [34]. Moreover, TIMM44 is positioned to act as a sensor for mitochondrial stress because the protein translocation motor is sensitive to ATP levels and misfolded proteins that accumulate in the matrix may subsequently impair protein import [5].

Alternatively, Arany and colleagues have shown that the adenine nucleotide transporter (ANT1 and ANT2) are linked to the TIM23 translocation channel by TIMM44 and these interactions may impact mitophagy [35]. ANT is proposed to act as a stress sensor that indirectly inhibits the TIM23 translocon by interaction with TIMM44. Under stress conditions, ANT can inhibit protein import, stabilize PINK1 on the OM, and induce mitophagy. Thus, MB-10-bound TIMM44 may have impaired interactions with ANT and mitophagy may be subsequently blocked. In sum, published studies indicate that TIM44 may play a central role in PINK1 trafficking at the TIM23 translocon during stress and support that future mechanistic studies with MB-10 are warranted.

### Small molecules as rapid modulators for regulating mitophagy

Small molecule modulators for mitochondrial protein import can be used to characterize the PINK1 import pathway and selectively modulate mitophagy in cell models. Previous approaches have relied on RNAi or CRISPR approaches, but with potential shortcomings. In the CRISPR approach, the target protein, such as TOMM7, can be deleted and cell lines can be established [23]. However, cells may have general impairment in mitochondrial protein import and may also adapt over time to overcome loss of the specific target protein. Alternatively, RNAi approaches may lead to secondary defects in protein import, particularly if a protein is essential, such as TIMM23 [23]. Under these conditions, general mitochondrial function is slowly impaired as the protein is removed. In addition, deletion of the candidate protein is not necessarily the same as inhibiting the protein with small molecules. Indeed, RNAi approaches to remove TIMM44 did not impact PINK1 trafficking in the same way that MB-10 altered PINK1 trafficking. This supports the use of small molecules to rapidly modify TIMM44 function. Of course, small molecules can also have off target effects, but the use of SAR compounds, such as Analog-3 and −4 that did not alter PINK1 trafficking, allow for assigning specific functions to MB-10 in modulation of TIMM44.

In sum, the use of MB-10 as a probe indicates that TIMM44 has a critical role in PINK1 trafficking. Future studies with MB-10 are geared to focus on molecular mechanisms of PINK1 interaction in the TIM23 translocon. Long-term, probes such as MB-10 will be fruitful for regulating mitophagy and for understanding how manipulation of mitochondrial protein import is a potential therapeutic platform for regulating mitochondrial stress and mitophagy.

## Experimental procedures

### Cell culture and transfection methods

HeLa cells including those with stable expression of EGFP-Parkin [8] were cultured in Dulbecco’s modified eagle medium, 25 mM glucose, 1 mM pyruvate (Life Technologies 11995-065), supplemented with heat-inactivated 10% FBS (v/v; Atlanta Biologicals S11150H) and 1% penicillin/streptomycin (Life Technologies 15140122) in a humidified incubator at 37° C under 5% CO_2_ (v/v). Tranfection was performed with jetPRIME transfection reagent (Polyplus 114-07) according to the manufacturer’s instructions. TIMM44 and negative control siRNA (QIAGEN SI03125969 and 1022076) were transfected at 20 nM using jetPRIME reagent (Polyplus 114-07) according to manufacturer’s instructions.

### Small molecule treatment with cultured cells

Small molecules are dissolved in 100% DMSO in brown glass vials (Wheaton 224750) and stored in a dessicator at room temperature. The MB-10 stock was 20 mM; fresh stocks were made monthly. Cells in culture were grown to 100% confluency followed by seeding to 25% confluency. After 48 hrs, cells were treated with MB-10 dissolved in fresh media; the final DMSO for all conditions was 1%, unless otherwise noted. For all experiments analyzed by BN-PAGE, 10 cm dishes (CELLSTAR 664-160) were used with 10 mL of media; all other experiments used 6 cm dishes (CELLSTAR 628-160) with 2.5 mL of media. At the time points indicated in each figure, media was aspirated and the cells were washed with PBS.

### Mitochondrial purification for immunoblotting analysis

Cells were removed from the culture dishes by incubating in Trypsin-EDTA (0.05%, Life Technologies 25300-054) for 5 min at 37°C. Cells were collected in media and centrifuged at 770 x *g* for 5 min. at 4°C. Cells were washed with ice-cold PBS (Life Technologies, 14190-144) and centrifuged again. The pellet was resuspended in 1 ml homogenizer buffer (20 mM HEPES-KOH pH 7.4, 220 mM mannitol, 70 mM sucrose) supplemented with 2 mg/mL BSA and 0.5 mM PMSF. Cells were lysed on ice by 20 strokes through a 25-gauge x 5/8 needle (Becton Dickson 305122) and 1 mL syringe followed by centrifugation at 770 x *g* for 5 min. at 4°C to remove unbroken cells. The supernatant was centrifuged at 10,000 x *g* for 10 min at 4°C, washed with homogenizer buffer (lacking BSA or PMSF), and re-centrifuged at 10,000 x *g* for 10 min at 4°C to obtain mitochondrial pellets. The pellets were resuspended in 100 μl of homogenizer buffer, and the protein concentration was determined using the Pierce BCA Protein Assay Kit (Thermofisher). For each condition, mitochondria (20 μg) were aliquoted into tubes and centrifuged at 10,000 x *g* for 10 min at 4°C. The supernatant was removed and mitochondria were disrupted with Laemmli-SDS sample buffer; the proteins were separated by SDS-PAGE followed by transfer to PVDF membranes, and analyzed by immunoblotting for indicated proteins.

### Antibodies

The antibodies used in this study include anti-Mortalin (UC Davis/NIH Neuromab Facility clone N52A/42, 75-127), anti-Parkin (Santa Cruz Biotechnology, sc-32283), anti-AIF (Santa Cruz Biotechnology, sc-13116), anti-TIMM23 (BD Biosciences, 611223), anti-PINK1 (Novus Biologicals clone 8E10.1D6, NBP2-36488), anti-TOMM20 (Santa Cruz Biotechnology, sc-11415), anti-TIMM44 (Proteintech, 13859-1-AP), anti-TOMM70 (Proteintech, 14528-1-AP), and anti-LRP-130 (Santa Cruz Biotechnology, sc-66844). The GST-MFN-1 (gift from Richard Youle), anti-PreP, anti-YME1L, anti-GFP, and anti-TOMM40 were polyclonal antibodies generated by vendor Pacific Immunology from recombinant proteins.

### Mitochondrial purification and in vitro protein import assays

Mitochondria isolation and imports were based on established methods [22]. [^35^S]-labeled proteins were synthesized in vitro via a coupled transcription/translation rabbit reticulocyte kit (Promega) incubated at 30°C for 90 min with [^35^S]Met/Cys metabolic labeling reagent (MP Biomedical, 015100907) and plasmid DNA expressing human PINK1, CHCHD2, or Su9-DHFR via an Sp6 promoter or MFN1-myc via a T7 promoter.

Isolated mitochondria were diluted to 0.4 μg/μL in homogenizer buffer (70 μg per reaction) supplemented with 17.5 μL of 10X energy generating solution (10 mM ATP, 5 mM magnesium chloride, 200 mM sodium succinate, 50 mM NADH in homogenizer buffer) and 0.1% BSA adjusted to a volume of 140 μL. DMSO vehicle or compounds dissolved in DMSO (1% final) were added to the mitochondria and incubated at 25 °C for 15 min. Reactions were then initiated by addition of 35 uL of precursor and aliquots were withdawn and stopped at specified time points.

The import of Su9-DHFR was terminated by the addition of ice-cold 10 μg/mL trypsin (Sigma) in homogenizer buffer. After 15 min on ice, soybean trypsin inhibitor (VWR Amresco) was added at 50 μg/mL. Mitochondria are pelleted by centrifugation at 10,000 x *g* for 5 min at 4°C. Import of OM proteins (PINK1 and MFN1-myc) was terminated by the addition of ice-cold homogenizer buffer and subsequent centrifugation at 10,000 x *g* for 5 min. Samples were then subject to carbonate extraction in 100 mM Na_2_CO_3_ at pH 11 [36]. For all precursors, after final recovery by centrifugation, mitochondria were disrupted in Laemmli SDS-sample buffer and analyzed by SDS-PAGE and autoradiography.

### BN-PAGE Analysis

After import of radiolabeled proteins into mitochondria (40 μg per reaction), reactions were stopped by the addition of ice-cold homogenizer buffer. Samples were centrifuged at 10,000 x *g* for 5 min at 4° C and gently washed with homogenizer buffer. The mitochondrial pellets were dissolved in ice-cold lysis buffer [20 mM HEPES-KOH pH 7.4, 50 mM NaCl, 1% digitonin (Gold Biotechnology D-180-2.5), 10% glycerol, 2.5 mM MgCl_2_, 0.5 mM PMSF, 0.5 mM EDTA] to give 2.5 μg of mitochondria per % digitonin and kept on ice for 15 min. Samples were subsequently centrifuged at 20,000 x *g* for 5 min at 4°C. Supernatants were then separated by BN-PAGE. Once complete, the gels were cut into two sections. One section was analyzed by immunoblotting and the second section was analyzed by a phospho-screen and imaged with a Bio-Rad Molecular Imager FX Pro Plus.

### Sub-mitochondrial localization

Mitochondria were isolated from cells using a Teflon dounce. Mitochondria (50 μg) were subsequently incubated in homogenizer buffer (Mitos), 20 mM HEPES-KOH pH7.4 (to generate mitoplasts), or 1% Triton X-100 (TX-100) supplemented with or without 3 μg/mL proteinase K. Samples were incubated on ice for 30 min, followed by inhibition of proteinase K with 1 mM PMSF. The samples were centrifuged at 20,800 x *g* for 10 min at 4°C. The supernatant was precipitated in 10% trichloroacetic acid (TCA, w/v) followed by centrifugation to recover the precipitate. The samples were disrupted in Laemmli-SDS sample buffer, separated by SDS-PAGE, and analyzed by immunoblotting.

### Carbonate extraction

Isolated mitochondria (50 μg) were incubated in homogenizer buffer or 100 mM Na_2_CO_3_ at increasing pH for 30 min on ice. Samples were centrifuged at 20,800 x *g* for 30 min at 4°C. The supernatant was TCA-precipitated and pellet and supernatant samples were disrupted in Laemmli-SDS sample buffer, separated by SDS-PAGE, and analyzed by immunoblotting.

### Immunofluorescence microscopy

Thirty min prior to fixing, MitoTracker Red CMXRos (Invitrogen, M7512) was added to each well to 20 nM final concentration. Cells on D-poly-Lysine coated coverslips were then fixed with pre-warmed 3.7% formaldehyde (v/v; VWR, VW3408-1), permeabilized with ice-cold methanol, and blocked with 1% fatty acid free BSA in PBS (v/v, PBS-BSA).

Where indicated, cells were subsequently incubated overnight at 4°C with indicated primary antibodies diluted in PBS-BSA. Following thorough washing in PBS, cells were incubated for 1 hr at room temperature with Alexa Fluor 350-conjugated secondary antibodies (Life Technologies) diluted in PBS-BSA. The following antibodies were used: anti-Mortalin primary with Alexa Fluor 350 goat-anti-mouse IgG (A11045), anti-TOMM20 primary with Alexa Fluor 350 goat-anti-rabbit IgG (A11046), and anti-DNA (PROGEN Biotechnik clone AC-30-10, 61014) mouse monoclonal IgM with Alexa Fluor 350 goat-anti-mouse IgM (A31552). Cells were then set on slides and images were obtained using a Leica TCS SPE DMI 4000B inverted confocal microscope.

### Mitophagy analysis

Mouse embryonic fibroblasts stably expressing Flag-Parkin [31] were transiently transfected with a RFP-GFP-LC3 plasmid using Polyplus jetPRIME^®^ transfection reagent according to the manufacturer’s instructions. Cells were allowed to incubate for 16 hr and then split into 6-well plates containing a sterile 18 mm coverslips. Analysis was performed in triplicate for each condition. Cells were incubated for an additional 24 h prior to MB-10 treatment. Cells were pre-treated with 1% DMSO or 40 μM MB-10 for 2 hours followed by the addition of 20 μM CCCP. At time 0 followed by 1, 2, and 3 hours, cells were washed twice with PBS and fixed in 4% paraformaldehyde in PBS for 15 minutes. Cells were subsequently washed twice with PBS and mounted using VECTASHIELD Mounting Medium with Dapi. Images were taken with a 63X oil objective Leica fluorescence microscope. 10 fields per coverslip with a total of 50 cells per condition were quantified. Green and yellow particles were quantified using color thresholding with a minimum particle size of 0.15 microns^2^. Red particles were quantified in a similar manner with the same minimum particle size.

### qPCR analysis

RNA was isolated from cells using TRIzol reagent, residual genomic DNA was removed with TURBO DNA-free kit, and cDNA was synthesized from the RNA with the Superscript III First-Strand Synthesis System (ThermoFisher 15596026, AM1907, and 18080051) according to manufacturer’s instructions. The qPCR was performed with 2x SYBR Green qPCR Master Mix (Bimake B21202) on the Biorad CFX Connect Real-Time PCR detection system. *PINK1* and *HRPT1* primers were obtained from Integrated DNA Technologies. *PINK1* primer sequences used were 5’-GGA CGC TGT TCC TCG TTA-3’ and 5’-ATC TGC GAT CAC CAG CCA-3’, as previously described [37].

### Statistical analyses

Data was plotted using GraphPad Prism version 5.02. Statistical significance of observations was determined using unpaired t tests (two-tailed), unless otherwise noted.

## Data availability

All data are contained within the article.

## Acknowledgments

We thank Dr. Janos Steffen and Dr. Juwina Wijaya for technical assistance as well as Dr. Richard Youle for the EGFP-Parkin HeLa cell line and the GST-MFN-1 antibody and Dr. Lars Dreier for FLAG-Parkin MEF cell lines.

## Funding and additional information

This work was supported by National Institutes of Health Grants DK101780 and GM61721, California Institute of Regenerative Medicine Grants RT307678, and USAF-AFRL FA9550-15-1-0406 (to C. M. K.); U.S. Public Health Service NRSA Grant T32GM07185 (to M.A.C.). The authors declare no competing financial interests.

## Abbreviations

CCCP: carbonyl cyanide 3-chlorophenylhydrazone
IM: inner membrane
IMS: intermembrane space
MitoBloCK/MB: <u>Mito</u>chondrial protein import <u>blo</u>ckers from the <u>C</u>arla <u>K</u>oehler laboratory
MTS: mitochondrial targeting sequence
OM: outer membrane
OMS: outer mitochondrial membrane localization signal
PAM: protein associated motor
PINK1: PTEN-induced putative kinase 1
SAR: structure-activity relationship
TIM: translocase of inner membrane
TOM: translocase of outer membrane

**This article contains supporting information**

