## Supplemental Figures for "A small molecule probe elucidates the role of mitochondrial translocase TIMM44 in PINK1/Parkin regulated mitophagy"

**Supplementary Information**

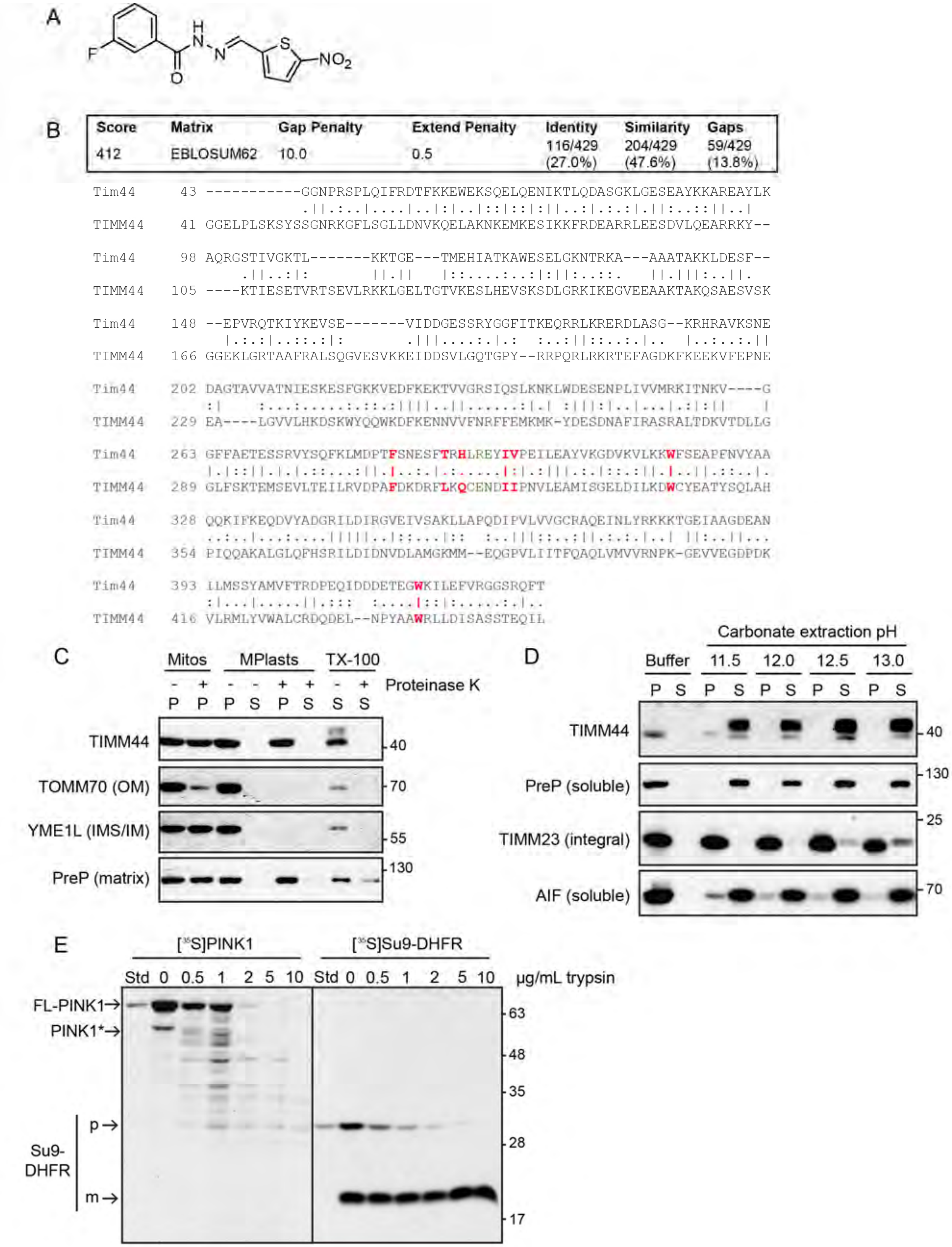


### Supplementary Figure 1

**Supplementary Figure S1.** Human and yeast Tim44 are highly conserved. **A.** MB-10 structure. **B.** A sequence comparison between yeast Tim44 and human TIMM44 (lacking the MTS). Information about the conservation score is provided above the sequences; single dot = dissimilar residues, two dots = similar residues, straight line = identical residues. The 7 residues that were identified as important for MB-10 binding [27] are highlighted in red. **C.** Isolated HeLa mitochondria were incubated in normal buffer (Mitos), hypotonic buffer to rupture the OM and generate mitoplasts (MPlasts), or Triton X-100 (TX-100) for lysis in the presence and absence of proteinase K. The pellet (P) was separated from the supernatant (S) and fractionated by SDS-PAGE. The indicated proteins were detected by immunoblotting. Controls include presequence peptidase (PreP) for the matrix, YME1L for the IMS, and TOMM70 for the OM. **D.** Isolated HeLa mitochondria were incubated in iso-osmotic buffer or 100 mM $Na_2CO_3$ at the indicated pH. Samples were centrifuged to separate the soluble (S) proteins from integral membrane (P) proteins in the pellet; markers include PreP (soluble), TIMM23 (integral), and AIF (soluble). **E.** Radiolabeled PINK1 or Su9-DHFR was imported into isolated HeLa mitochondria. Post-import, the samples were treated with the indicated concentration of trypsin on ice for 15 min before separation by SDS-PAGE. Abbreviations, p, precursor and m, mature forms of Su9-DHFR; FL-PINK1 for full-length PINK1 and PINK* marking the PARL-cleaved form.

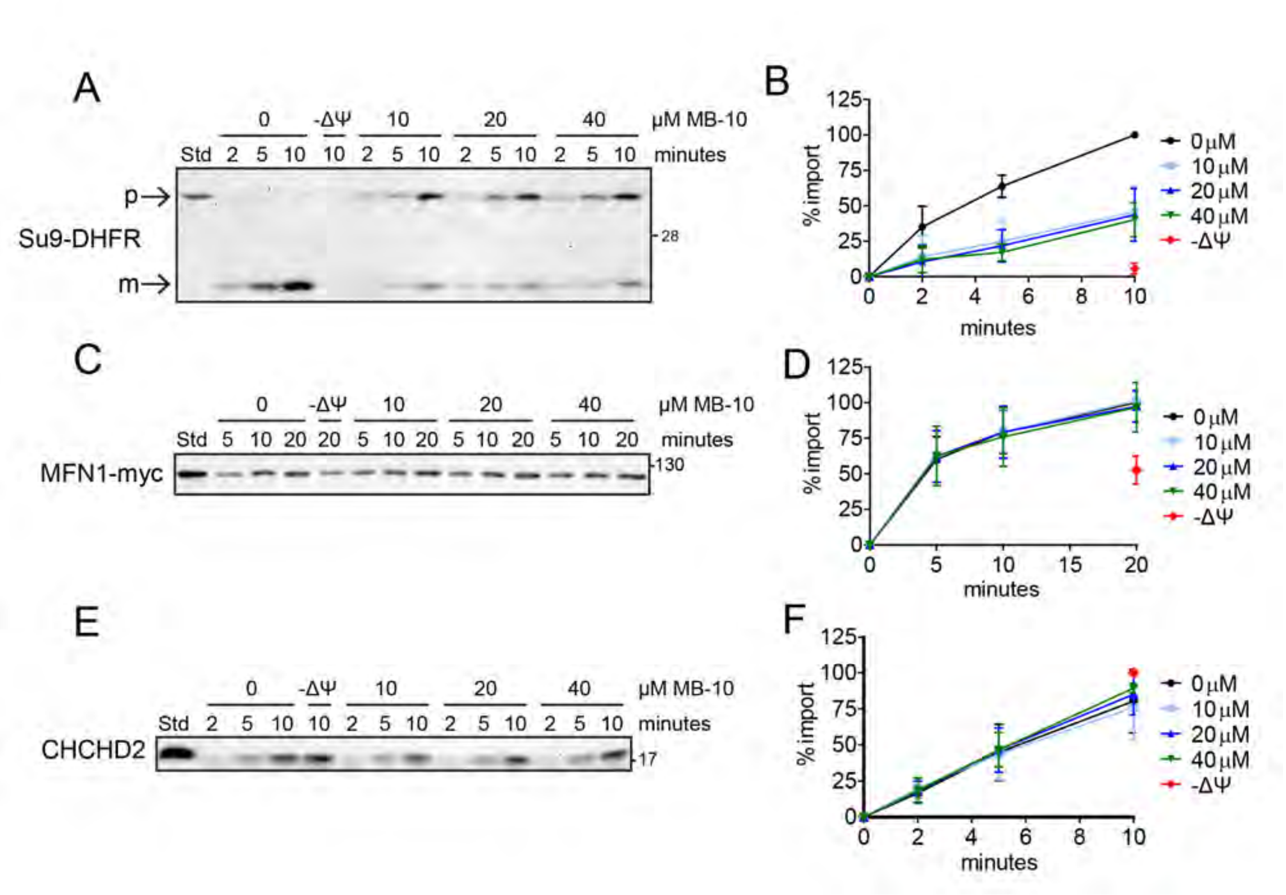


#### Supplementary Figure 2

**Supplementary Figure S2.** MB-10 inhibits the import of precursors that use the TIM23 import pathway. **A.,C.,E.** Representative images of time course imports for radiolabeled precursors into isolated HeLa mitochondria after a 15 min incubation with MB-10 or DMSO; -Δψ = 50 μM CCCP. Precursors included Su9-DHFR (matrix), MFN1-myc (OM), and CHCHD2 (IMS). **B., D., F.** Quantification of the imports from '**A**', '**C**', and '**E**'; data represent the average ± SD of *n* = 3 trials. In '**A**', the 'm' band was quantified. The longest time point at 0 μM MB-10 band represents 100% import for '**B**' and '**D**', and the -Δψ band represents 100% for '**F**'. p, precursor and m, mature.

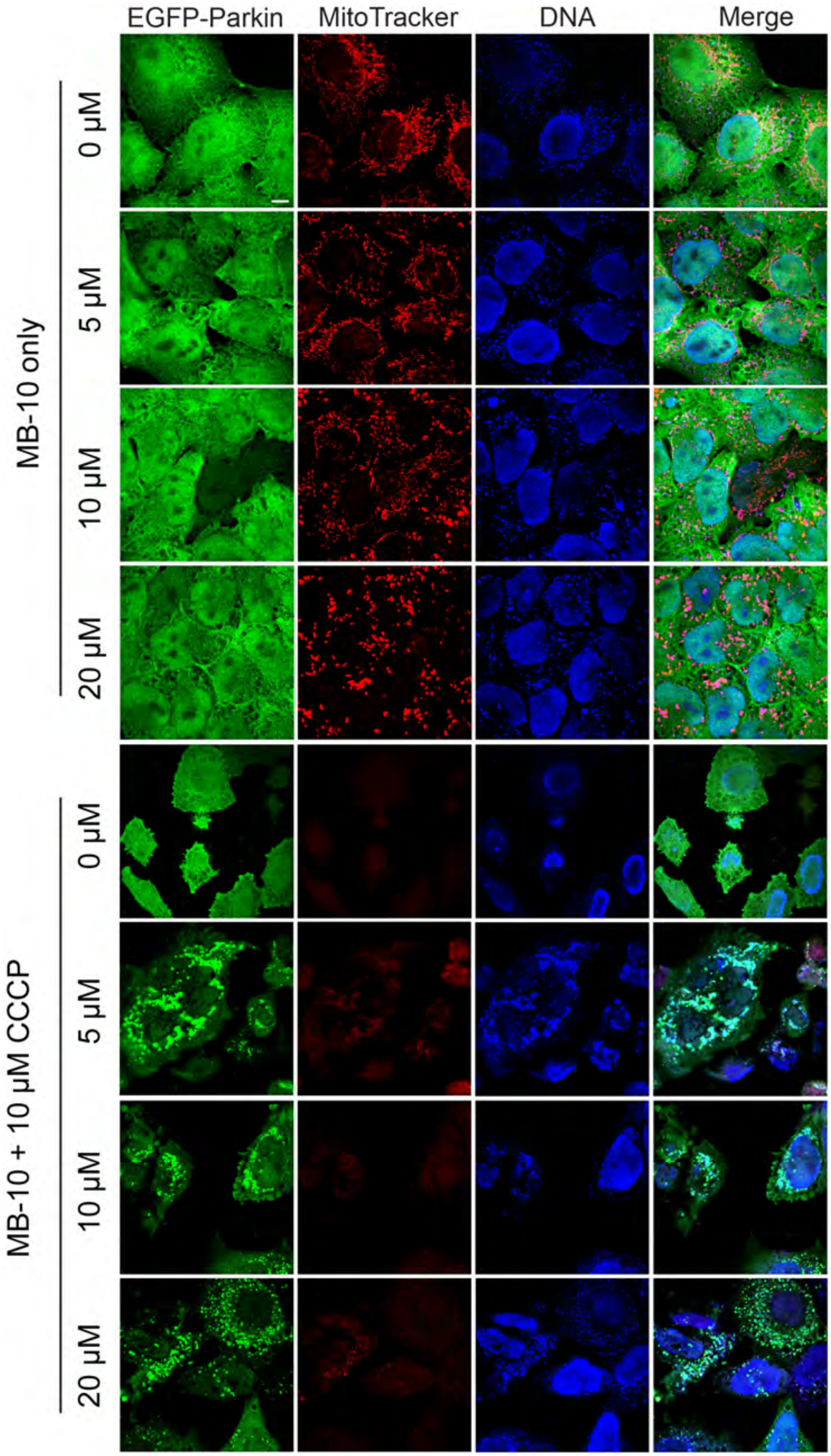


**Supplementary Figure 3**

**Supplementary Figure S3.** MB-10 inhibits mitochondrial DNA degradation in CCCP-induced mitophagy. Representative images of HeLa cells with stable overexpression of EGFP-Parkin after treatment with the indicated concentration of MB-10 for 2 hrs, followed by the addition of either DMSO of 10 μM CCCP for an additional 24 hrs. Both nuclear and mitochondrial DNA (as cytosolic puncta representative of nucleoids) were stained with an anti-DNA antibody. MitoTracker Red marks mitochondria with a membrane potential. Scale bar = 7 μm.
